# Genetic patterning in Central Eurasia: population history and pigmentation

**DOI:** 10.1101/255117

**Authors:** Ellen C. Røyrvik, Nadira Yuldasheva, Susan Tonks, Bruce Winney, Ruslan Ruzibakiev, R. Spencer Wells, Walter F. Bodmer

## Abstract

Central-western Asia has often been underrepresented in population genetic studies, but it is important for the clarification of the peopling of Eurasia and the relationship between its western and eastern extremities. We genotyped individuals from over 40 population groups, mostly central Eurasian, for mitochondrial HVR1, *CCR5del32* and five functional *MC1R* variants (p.Val60Leu, p.Val92Met, p.Arg151Cys, p.Arg160Trp, p.Arg163Gln), and collected published genotype data for comparison. Mitochondrial profiles confirm both the higher heterozygosity in Central Asia than in surrounding areas, and the broadly northern European distribution of *CCR5del32*. The *MC1R* variants profile alone is a good determinant of the longitudinal position of a population group, and combined F_*ST*_ values divide Eurasia into seven broad geographic divisions. We can conclude that Central Asia shares genetic features with both eastern and western Eurasia, compatible with both a scenario where the former acted as a source for the latter two’s genetic diversity, or one where Central Asia is a ‘hybrid zone’ where eastern and western peoples met. Furthermore, the overall high F_*ST*_ values for functional *MC1R* variants combined with presumed selection pressures on skin pigmentation in low-UV areas lead us to conclude that different variants were selected for in east and west Eurasia, an example of convergent evolution.

## Introduction

Western and Central Asia, occupying a vast and pivotal area of Eurasia, have a unique place in the continent’s population history. Many migration waves, from prehistoric to early modern times, have started, ended or passed through these areas, possibly changing biometrical phenotypes, language, and customs [1, 2]. However, the genetic profiling of its human populations has mostly lagged behind that of Europe and East Asia. The two central regions are interesting from two aspects. Western Asia, perhaps Central Asia as well, is considered to be the source area, after Africa, for population expansion into Europe and East Asia, so it is important for putting the latter areas in context. It is also, in its own right, important for the elucidation of global human genetic variation. In this paper we examine several markers that have substantial known allele or haplogroup frequency differences between Europe and East Asia, in a large set of populations from across Eurasia, with a particular focus on Central and Western Asia. The markers examined are five functional coding SNPs in *MC1R*, associated with fair complexion and red hair (rs1805005, rs2228479, rs1805007, rs1805008, and rs885479), the 32 base pair deletion in *CCR5* (rs333) and the mitochondrial hypervariable region 1. We also relate these data to previously published Y chromosomal data from the same populations [3]. The Y chromosome and mitochondrial data effectively serve as controls for population characterisation in the search for differential effects, particularly selection associated with the *MC1R* and *CCR5* variation.

*CCR5*, a member of the seven-pass transmembrane G-protein coupled receptor family [4], is a chemokine receptor notable for being a co-receptor for viruses, including, in particular, HIV. A *CCR5* variant containing a 32 base pair deletion coding for a non-functional receptor has reached surprisingly high frequencies in some populations, possibly due to a role in limiting viral or other infections [5, 6]. In this paper we extend the data on the striking geographic distribution of *CCR5del32* [7].

*MC1R* was the first gene whose variation was shown to have a demonstrable role in human pigmentation variation [8], having variants that cause red hair, pale skin and poor tanning ability in northwestern Europeans. Encoded by a single exon, the coding region is small and has unusually high nucleotide diversity [9]. The *MC1R* variants were investigated as Eurasia hosts some of the most extreme differences in pigmentation in the world, particularly with respect to hair color. Diversity in external pigmentation is one of the most visually striking differences between human ethnic groups. Basal external pigmentation is largely genetically determined, and there is substantial geographical patterning in the frequencies of alleles of genes involved particularly in melanin production and distribution [10]. As pigmentation variation largely correlates with distance from the equator and UVR intensity, it is most obviously related to natural selection associated with climatic differences [11, 12]. Inevitably, variation in the distribution of pigmentation gene variants may also be influenced by the population histories of the areas in question.

## Materials and Methods

Samples novel to this study were collected and treated as described in [3] and [13], all with relevant ethical permissions. Additional allele and genotype data from the literature were collected for *MC1R* variants and *CCR5del32*. In all cases, samples were from unrelated individuals not subject to any known relevant medical conditions (e.g. melanoma or AIDS). See S1 Table and S2 Table, respectively, for a synopsis of the samples genotyped for *MC1R* and *CCR5*, and Fig 1 and S1 Fig for maps of sampling locations. A subset of the 746 samples described in [3] was typed for mitochondrial hypervariable region 1 (HVR1). Some of these samples have also been typed for mitochondrial DNA (mtDNA) in [14, 15]. In total, genetic information on 58 855 individuals was collected. The geographic locations for these samples span all of Eurasia, including areas that are often poorly covered, such as Central Asia and southern Siberia. See S1 Appendix for additional detail on included samples.

**Fig 1.**
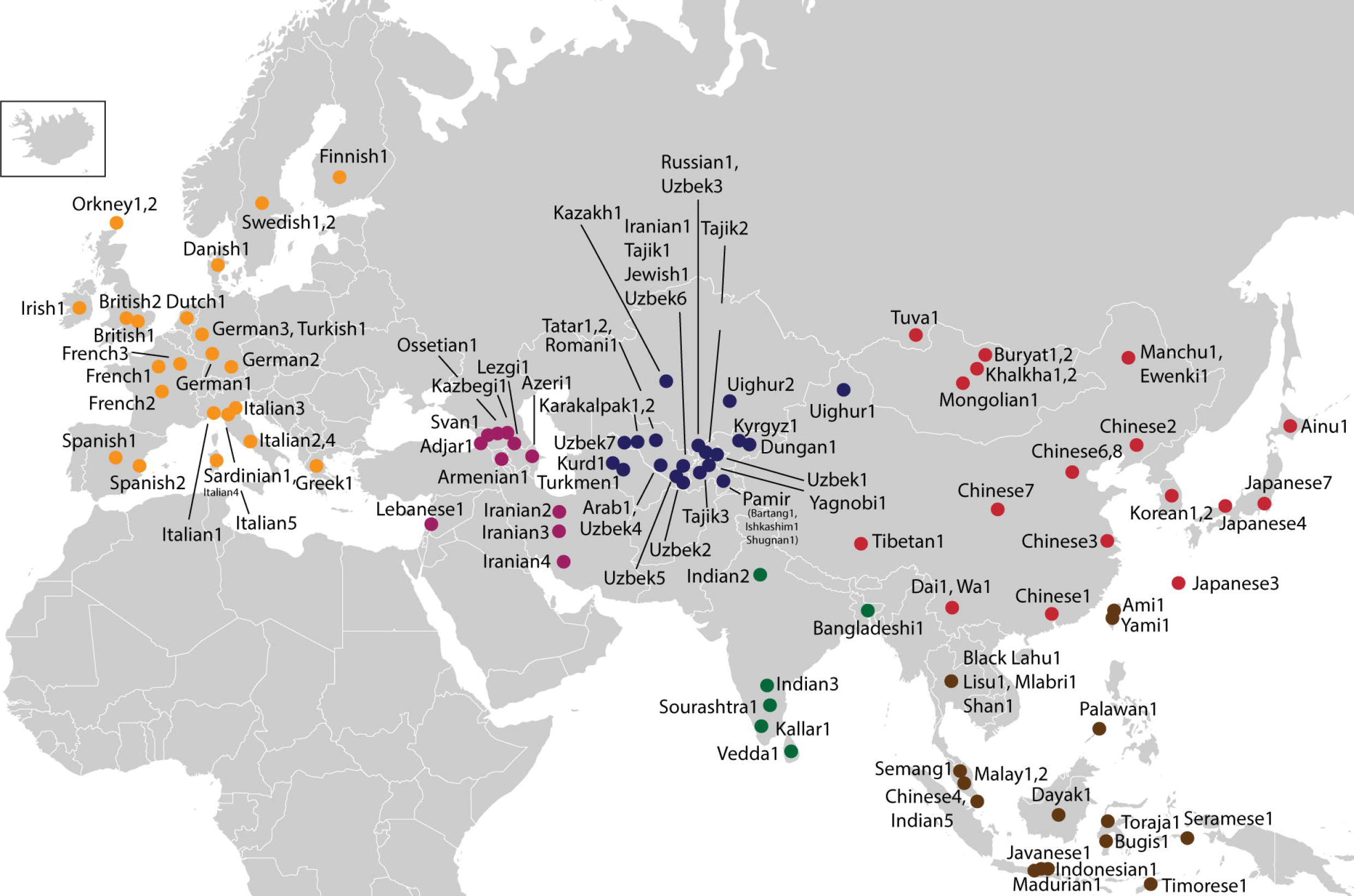
Sampling locations for individuals genotyped for *MC1R* variants. Locations are colored according to region, with Europe - orange, West Asia - purple, Central Asia - blue, South Asia - green, East Asia - red, and Southeast Asia - brown.

Genotyping for *MC1R* variants (p.Val60Leu – rs1805005, p.Val92Met – rs2228479, p.Arg151Cys – rs1805007, p.Arg160Trp – rs1805008, and p.Arg163Gln – rs885479) and the *CCR5* 32bp deletion was performed using PCR, and products were resolved by gel electrophoresis or alkaline-mediated differential interaction (AMDI) [16]. HVR1 was sequenced between nucleotide positions 16001 and 16571 of the revised Cambridge Reference Sequence 17. Haplogroups were assigned according to published data (see S2 Appendix for references). All primer sequences and PCR conditions are available on request.

F_*ST*_ for all variants was calculated using Arlequin 3.11 [17]. Regression analyses for the *MC1R* variants, mitochondrial heterozgosity, and the *CCR5* deletion were performed with default linear models, and Pearson’s correlation coefficient for mitochondrial and Y heterozygosity was calculated using R v.2.9.2. Principal components analysis was performed in R v.2.9.2 using default settings. Partial Mantel tests for language-genetics correlation were performed using the R package vegan, using F_*ST*_ for genetic distance, great circle geographic distances, and a heuristic distance scheme for languages based on language family affiliation (dialectal variants were given a distance score of 0.5, members of same sub-family were scored as 2, members of same family as 3, and unrelated languages as 1000), all with 999 permutations. For a Mantel test examining residual correlations between MC1R variant F_*ST*_s and UV exposure after controlling for geographical distance, exposure was calculated by taking the average of daily erythemal UV measurements between August 1996 and August 2003, with raw data files downloaded from the NASA TOMS (Total Ozone Mapping Spectrometer) website, *http://ozoneaq.gsfc.nasa.gov*.

## Results

Variant frequencies for *MC1R* and the *CCR5* deletion are shown in S3 Table and S4 Table, and the mitochondrial haplogroup frequencies in S5 Table.

### CCR5del32

The deletion reaches its highest frequencies in the circum-Baltic area, with the Byeolrussian sample having a frequency of 16.25%, and its lowest in East and Southeast Asia, with no carriers in the Mongolian, Taiwanese and Burmese samples, and only 16 among 3702 Chinese chromosomes (0.43%). The frequency of the deletion decreases approximately radially from the eastern Baltic (see Fig 2). F_*ST*_ shows significant differences between Northwest Eurasians and populations from East Asia, and between Northwest Europeans and populations from Southern Europe (see S6-8 Table).

**Fig 2.**
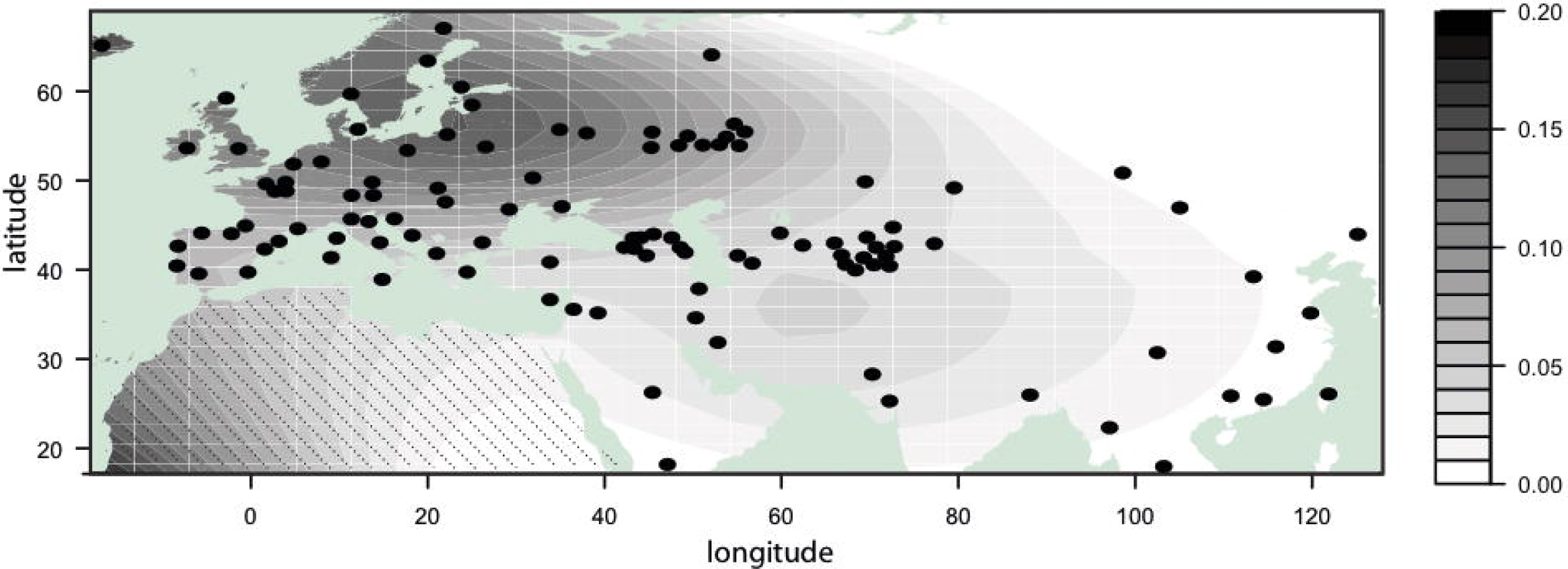
Frequency distribution of *CCR5del32*. The darker color indicates higher frequencies, according to the bar on the right, and the contour lines separate zones of different frequencies. Sampling locations are marked by black dots. Note that no samples were sourced from Africa.

### Mitochondrial DNA

The haplogroup frequencies observed are in line with what has been previously observed for the study region [14, 15, 18]. Populations of known higher Turkic input (Kyrgyz, Kazakhs, Karakalpaks) have frequency profiles displaying more typically East Asian haplogroups (A, B, C, D, M*), populations from the Caucasus principally have haplogroups associated with western Eurasia, and sedentary Uzbek and Tajik profiles are intermediate (see Fig 3). The Iranians from Uzbekistan likewise have an intermediate profile, similar to their Tajik and Uzbek neighbours, and different from those living in Iran. F_*ST*_ values are low (0-0.073), but a broad differentiation between eastern and western parts of Eurasia may be discerned (see S6-8 Table). In terms of variability, the Central Asian populations have, with few exceptions, the highest heterozygosity measures (see Table 1). This is a similar, but stronger, pattern to what was seen in the Y chromosome data (Wells *et al*., 2001 [3]) for the same samples. The correlation between the two heterozygosity measures is 0.3, but statistically it is not significant.

**Fig 3.**
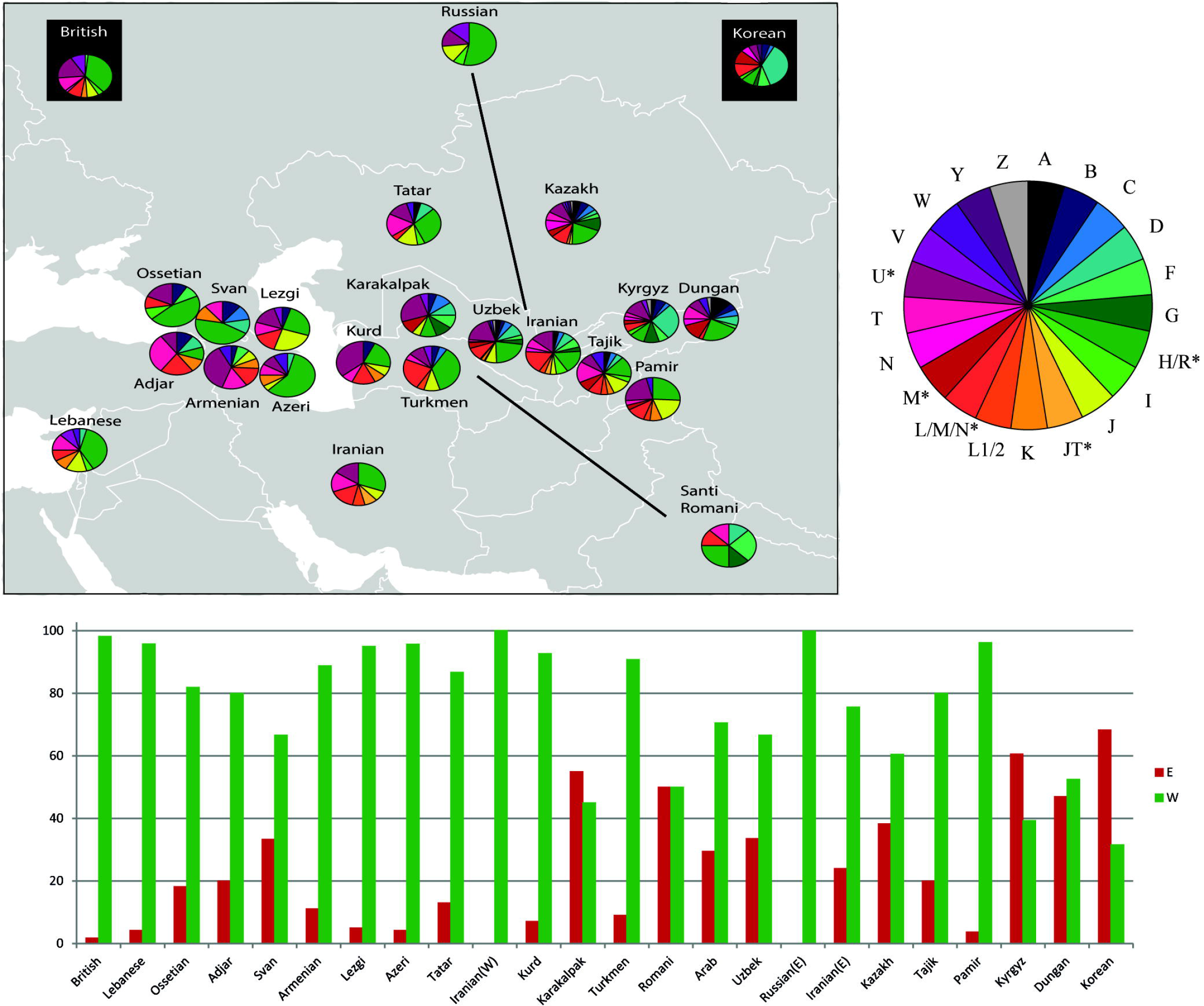
Mitochondrial haplogroup profiles in West and Central Asia. The wheel to the right gives the colour-haplogroup correspondences. The bar graph below gives the frequency per population of combined haplogroups, where green bars comprise haplogroups which are typically ‘West Eurasian’, and red bars the typically East Asian haplogroups.

**Table 1.** Mitochondrial heterozygosity. Populations ranked by increasing heterozygosity, defined as one minus the sum of squares of all the allele frequencies.

| Population | Heterozygosity |
| --- | --- |
| Azeri | 0.63 |
| Russian | 0.66 |
| Ossetian | 0.73 |
| Svan | 0.74 |
| Kurd | 0.79 |
| Turkmen | 0.79 |
| Armenian | 0.80 |
| British | 0.80 |
| Lebanese | 0.80 |
| Romani | 0.81 |
| Lezgi | 0.82 |
| Iranian(W) | 0.82 |
| Korean | 0.82 |
| Adjar | 0.82 |
| Pamir | 0.83 |
| Tatar | 0.83 |
| Arab | 0.86 |
| Karakalpak | 0.87 |
| Kyrgyz | 0.88 |
| Uzbek | 0.88 |
| Dungan | 0.89 |
| Iranian(E) | 0.89 |
| Tajik | 0.90 |
| Kazakh | 0.91 |

### MC1R

The five *MC1R* variants have substantially differing geographic distributions (see Fig 4 for a schematic overview). Broadly speaking, they fall into two groups: those that reach their highest frequencies in West Eurasia (p.Val60Leu, p.Arg151Cys, and p.Arg160Trp), and those that have their frequency peak in East Eurasia (p.Val92Met and especially p.Arg163Gln). For p.Val60Leu, the basal trend is of higher frequencies (*>*7%) west of the Caspian Sea, and lower frequencies (*<*5%) east of it. p.Val92Met has the most even distribution for most of the continent, but with higher levels following an arc from Southeast Asia to Northern Europe. p.Arg151Cys and p.Arg160Trp both reach their highest frequencies (up to 12-13%) in Northwest Europe, with levels decreasing in all directions, though p.Arg160Trp has more of an easterly spread showing higher frequencies in several population samples from the Caucasus than in Southern Europeans. p.Arg163Gln is most frequent in East Asia, and is the only *MC1R* variant under consideration to consistently reach frequencies of more than 50%. The Finns and Saami have, for Europe, uncharacteristically high p.Arg163Gln levels (30% and 15%, respectively, see S3 Table) which may be relevant to the origins of these populations. For all *MC1R* variants, frequencies show South Asians to be significantly different from all of the other populations we have studied (Fisher’s exact test, results not shown), in having very low frequencies for all the minor alleles we have typed for. In contrast, non-South Asian populations have relatively high frequencies for at least one of these alleles. This is consistent with these southern Asian populations not having been subject to pigmentation-reducing selection on the subcontinent due to the higher levels of UV-radiation there.

**Fig 4.**
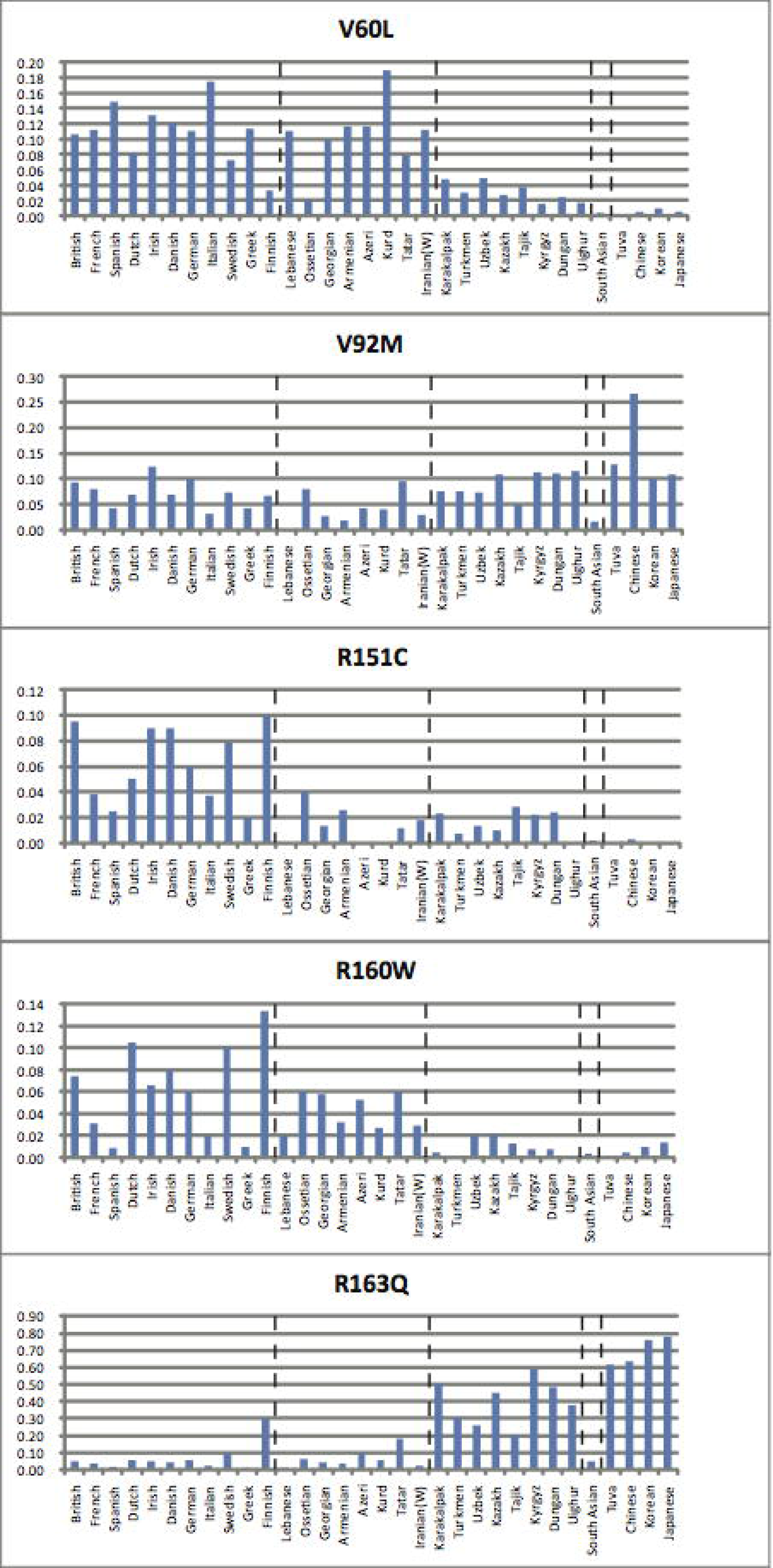
*MC1R* variant frequencies by longitude. Frequencies of the five *MC1R* variants for representative populations, plotted on a west-to-east (left to right) axis. Vertical dashed lines delineate geographical regions, from left to right: Europe, West Asia, Central Asia, South Asia and Eastern Asia.

F_*ST*_ measures reveal geographical distinctions along a longitudinal axis, excluding South Asia (see Fig 5, S6-8 Table and S9 Table). p.Arg151Cys and p.Arg160Trp distinguish Northern Europe from the rest of the continent. p.Val60Leu shows a tendency to separate Europe and West Asia from Central and East Asia. p.Arg163Gln, for which F_*ST*_ values between populations reach a maximum of 0.94, divides the continent into blocks approximately consisting of Europe and West Asia, Central Asia, and East and Southeast Asia. Central Asian populations that stand out are the Karakalpaks, Kyrgyz and Kazakhs of northern Central Asia, and the Mandarin-speaking Dungans from the border regions with China, all of which show a greater affinity with the East Asian block by more similar allele frequencies and lower F_*ST*_ values. The last variant examined, p.Val92Met, separates Southeast Asian and southern Chinese groups (Dai and Wa) from other populations. A principal components analysis of the *MC1R* variants confirms the above (see Fig 6), with PC1 (0.91 of the variance) separating East Asia from West Eurasia and South Asia, with Central Asia in a medial position. PC2 (0.04 of the variance) further separates Northern Europe from the rest of West Eurasia. PC1 largely mimics an east-to-west axis, probably principally due to the influence of p.Arg163Gln, while PC2, splitting the north of West Eurasia from the south, is likely influenced by the higher p.Arg151Cys and p.Arg160Trp levels in the north (see below).

**Fig 5.**
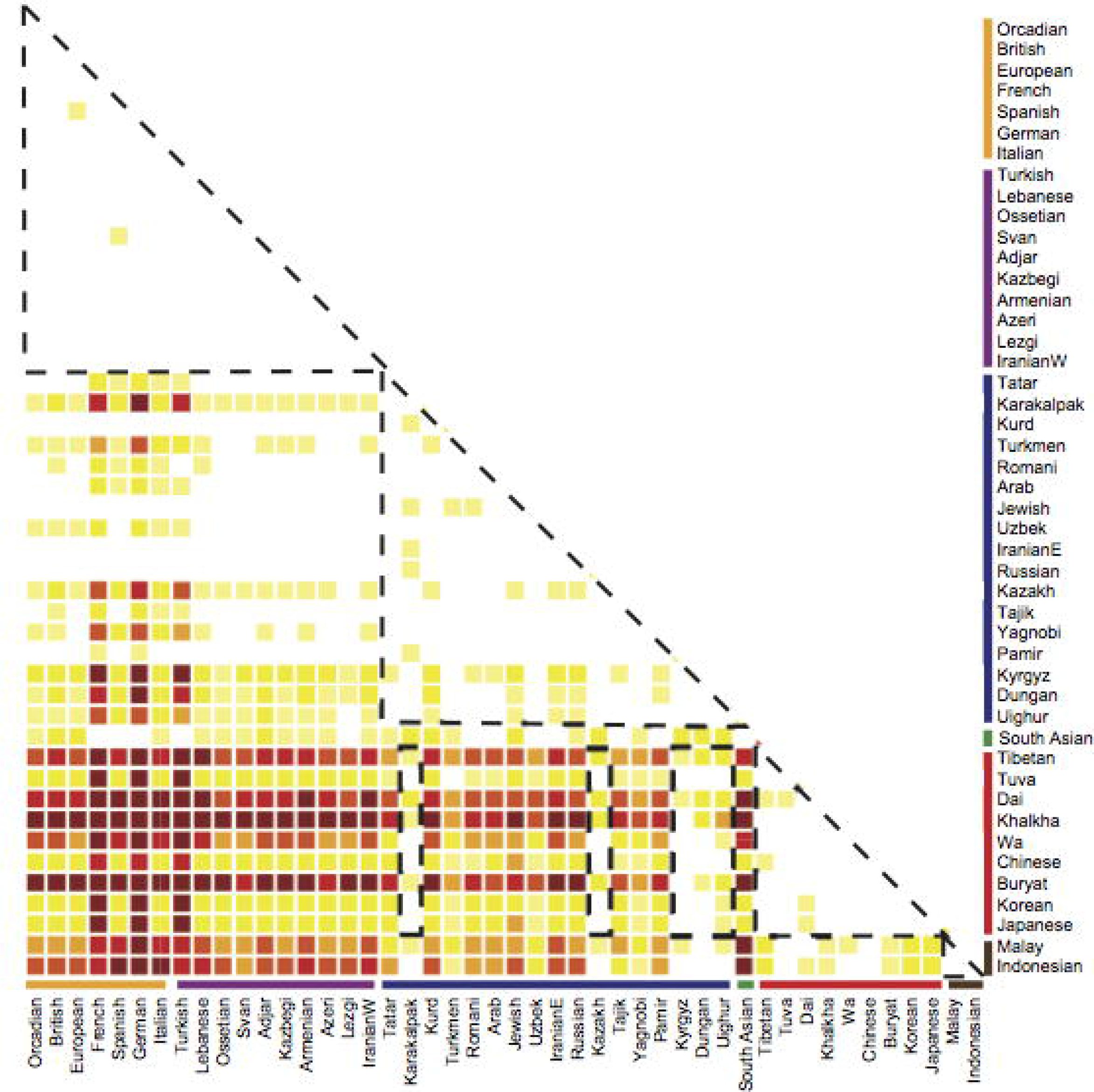
Graphical representation of *MC1R* variant F _*ST*_ values. Averages of separately obtained values for each variant, where white *<*0.05, pale yellow=0.05-0.1, yellow =0.1-0.2, pale orange=0.2-0.3, orange=0.3-0.4, red=0.4-0.5, dark red *>*0.5. Blocks of similar populations are indicated with dashed triangles along the diagonal. Dashed boxes, as an extension of the East Asian triangle, highlight the greater similarity of the Karakalpak, Kazakh, Kyrgyz, Dungan and Uighur groups to the East Asian populations than that shown by other Central Asian populations. Population names are colour-coded and arranged by continental region, see above and Fig 1.

One way to look at the variation in the frequencies of the *MC1R* variants is to consider the relationship of their frequencies with latitude and longitude. Within Europe p.Arg151Cys and p.Arg160Trp frequencies are good predictors of latitude, with r^2^ values of 0.72 for p.Arg151Cys and 0.88 for p.Arg160Trp, both being positive correlations (where latitude was regressed on frequency). As these two variants are the most strongly associated with pale skin [19], and UV-radiation decreases with increased latitude, this is not unexpected. Furthermore, the combined information provided by the frequency data for each of the *MC1R* variants in a population sample is a reasonable predictor of that sample’s longitudinal origin within the continent, with a multiple linear regression r^2^ = 0.69, where a linear combination of the frequencies is used as the predictor variable in the model and longitude is the response variable. A partial Mantel test was performed to see if there was any residual correlation between *MC1R* variant frequencies and erythemal UV exposure, after geographical distance was controlled for. No such correlations were observed for p.Val60Leu, p.Val92Met or p.Arg163Gln that were statistically significiant, but moderate and signficant correlations were found for p.Arg151Cys (Mantel r=0.47, p=0.001) and p.Arg160Trp (Mantel r=0.27, p=0.001).

## Discussion

### Population history

Central Eurasia holds a wealth of cultural and linguistic diversity, as well as genetic diversity. Central Asia’s earliest modern human inhabitants are deemed to have been Caucasoid by physical anthropologists, though their methodology has occasionally met with scepticism [20], and all its earliest languages on record (from the 6th century BC) belong to the Iranian branch of Indo-European. Nomadic pastoralists of an eastern Asian type (“Mongoloid”) appear in what is now Kyrgyzstan in the 5th-8th centuries AD, and this physical type is claimed to become dominant from the 13th century [1]. There are many historically attested waves of Turkic migration from the east into Central Asia, and across it to Western Asia, starting from around the 5th century AD, however, the number of individuals involved is difficult to estimate [2]. There are still remnants of people speaking the once-widespread East Iranian languages (e.g. the Pamirs), but most inhabitants now speak Turkic languages. Two of the largest ethnic groups in Central Asia, Uzbeks and Kazakhs, are recent (late 15th century) agglomerations under new names of former groups living in their current areas.

The above illustrates a very small portion of the genetic complexity one may expect to find in the region, though the general stability of the area’s population as a whole must be considered. Two main theories have been put forward as to the origins of Central Asia’s genetic character, which displays both typically European/West Asian and East Asian features. The first is that Central Asia has been a primary source of genetic variation for both Europe and East Asia, and the second that most currently observed genetic variants arrived with separate streams of migrants from Europe and East Asia, largely swamping the sparse extant population of the arid region. Different studies find cases for either hypothesis, mostly based on mitochondrial and Y chromosomal DNA markers [3, 14, 18, 21–23]. Mitochondrial DNA data tends to support a situation where Central Asia has been a contact zone for East and West Eurasian peoples, possibly outnumbering a dispersed population in the region that is, and was, primarily desert [14, 15, 24]. Indeed, Gonzalez-Ruiz *et al*. present ancient DNA evidence for that the area here defined as Central Asia was, from the mitochondrial point of view, western Eurasian until the Iron Age [25]. Arguments for this position include that there is little haplogroup overlap between East and West Eurasia (a smoother frequency gradient might be expected if they had all diffused from Central Asia), and that there is no great divergence of HVRI sequences between Central Asia and its neighbours to the east and west – that most haplotypes found in Central Asia are also found in Europe and East Asia – suggesting a more recent arrival [15]. A recent ADMIXTURE analysis using genome-wide, high-density SNP data can be interpreted in support of this, with Central Asian populations being described as unequal admixtures of typically Caucasian, South Asian, and East Asian genetic backgrounds [26]. However, it may be argued that frequency differences arising by drift in sparse, separated populations would not be expected to create such smooth gradients. A study of 54 794 SNPs in East and Southeast Asian populations concludes that the main genetic input for East Asian populations comes from Southeast Asia, with only a very small proportion from Central and South Asia [23]. Conversely, Y chromosome data are consistent with Central Asia being a source of pan-Eurasian genetic variation, detecting several emigratory events [3].

For the data presented here, *CCR5del32* is the least informative of general population history. The deletion is estimated to be around 5000 years old (3150-7800 years, 95% CI), has been found in Bronze Age skeletal remains [27, 28], and has most likely been subject to strong positive selection pressures [29, 30], though see Bollbäck *et al*. (2008) and Novembre and Han (2011) for a dissenting opinion [31, 32]. These selective pressures may be more intense in the northern part of the continent [30], as is suggested by the recent expansion of the deletion allele to high frequencies in otherwise quite dissimilar populations (e.g. Belgians and Tatars). However, this pattern could also be due to drift, once the allele was established; though as a frequency-dependent process, drift is stronger when frequencies are low.

The tendency observed in mitochondrial DNA for several Central Asian groups of ethnographically presumed high input from Turkic populations to have more East Asian haplogroups is mirrored in *MC1R*. For all *MC1R* SNPs, the F_*ST*_ values are consistently higher between more ‘Iranian’ groups (like the Pamirs, Tajiks, and Uzbeks) and Chinese (representing East Asia) than between more ‘Turkic’ groups (such as the Kazakhs, Karakalpaks, and Kyrgyz) and the Chinese: 0.08-0.10 versus 0.01-0.04. In addition, for the Y chromosome data from the same populations, Uzbeks, Tajiks, Pamirs and Turkmens display a higher percentage of ‘western’ haplogroups and Kazakhs and Karakalpaks have more haplogroups typical of Mongolia and China [3]. This pattern is likewise found in high-density SNP data focusing precisely on elucidating the geographic origin and expansion of Turkic speaking peoples [26].

The *MC1R* SNPs further reveal the notable property that, when the geographical structure from each of them is combined, they define geographic subdivisions of Eurasia quite well: Northern Europe, Southern Europe, West Asia, South Asia, Central Asia, East Asia, and Southeast Asia (see Fig 5 and S9 Table, in particular).

Though based on few loci, the data seem congruent with known historical incursions of Turkic peoples into, and occasionally through, Central Asia, overlaying a more Caucasoid (West Asian) character of prehistoric peoples in the region. Conversely, the general picture of higher genetic diversity in Central Asia compared with the rest of Eurasia may be interpreted as supporting a position for this region as a source of diversity for the east and west. A further demonstration that certain markers can be indicative of past population movements is seen in the uncharacteristically high frequency of p.Arg163Gln in Finns and Saami. If the variant can be seen as a marker of East Asian genetic influence, this eastern input is supported by a wide range of markers throughout the genome, from large SNP arrays as well as mitochondrial DNA, Y chromosome data, HLA haplotypes, and blood groups [33–37].

### Linguistic groups

The language groups that are most prevalent in West and Central Asia are Iranian and Turkic, represented in our samples by languages of both the western and eastern branches of Iranian, and languages from four different branches of the Turkic family (see S2 Fig for the relationships between these different languages, and their distribution within West and Central Asia) [1]. The populations speaking these languages were analysed separately for mitochondrial haplogroups and MC1R variants markers to see if there was any substructure in our genetic data that indicated affinities within a language grouping that was in contrast to the geographical locations of the samples. A limited amount of such correlation was found. p.Arg151Cys and p.Arg160Trp, being largely limited to Europe, showed no correlation with language in these mostly Central Asian populations. Partial Mantel tests controlling for geographic distance reveal close to statistically significant correlations between mitochondrial and p.Val60Leu F_*ST*_s and language affiliation (p = 0.07), and significant residual correlations were found between language affiliation and p.Val92Met/p.Arg163Gln F_*ST*_ values. The p.Arg163Gln correlation in particular is highly significant (p = 0.001 by permutation), where populations speaking Turkic generally have higher levels of p.Arg163Gln than geographically proximate Iranian-speaking groups. One publication, covering some of the same population groups, finds genetic structuring that is explained by linguistic affiliation [38]. However, their study, based on fewer populations and language groups and a smaller study area than the current one, is essentially comparing the genetics of ethnic groupings, rather than that of hierarchical linguistic relationships as represented by the speakers of language family subgroups. Nevertheless, all these results intimate that, while geographic distance remains a prime explanatory factor for genetic variation distribution, a genetic signature of population history, as represented by inherited language, may remain.

### Pigmentation evolution

Despite the demonstrated function of *MC1R* in integument pigmentation [8, 39–41], and that skin color is likely to be under considerable selective pressures [42], tests for selection in the gene have not provided clear-cut answers [9, 11, 41, 43–47]. Intuitively, as its product is one of the most influential elements in melanin production and distribution [19], *MC1R* is a prime candidate for selection to act upon in producing lighter skin tones in areas with lower sunlight, as is the case for other genes that affect pigmentation [10, 48]. Several studies also show that Europeans and East Asians owe their comparably light skin to different variants of pigmentation genes [10, 49]. Some assessments that did not find evidence for positive selection used methods unsuited to the probable selection pressures, both positive and negative. For example, Harding *et al*.’s [46] assessment depends on there having been no selection on *MC1R* in the course of evolution along the human and chimpanzee lineages. Norton *et al*. do, however, find evidence for selection at the locus, at least in Europe [10]. Two main characteristics of *MC1R* variation highlighted in this paper support the contention: The first is that of much higher levels of nucleotide variation in this very small gene (55 SNPs in 951bp) compared to a variety of other gene classes, which is a strong indication of selection [50]. The second is the geographical distribution of variation. In contrast to most loci, the majority of nucleotide diversity is found in Europe (44 SNPs), not Africa (11 SNPs). Furthermore, the European samples display a non-synonymous/synonymous substitution ratio of 4.5 (6 in northern Europeans), versus an African ratio of 0.37, which indicates adaptive selection in Europe and purifying selection in Africa. The derived alleles of the five *MC1R* SNPs assessed in the present study are all associated with a lighter skin tone [19, 51], and p.Val60Leu, p.Val92Met and p.Arg163Gln exhibit higher than expected differentiation between population groups as measured by F_*ST*_ (compared to subcontinental genomic averages from Norton *et al*. based on 11 078 autosomal SNPs [10]). The distributions of the variant alleles show a high degree of population differentiation, and this type of genetic structuring has been predicted and shown to be a feature of a sizeable proportion of selected variants [50, 52, 53]. The comparison of PCAs for the *MC1R* variants and the Y chromosome (see Fig 6) strengthens the case for selection at the former locus. The Y chromosome, while subject to accelerated drift compared to autosomal loci, is not subject to significant selection, and these analyses display very disparate patterns, for which selection is the most obvious causative factor. A further indication of the importance of *MC1R* variants as targets of selection is the discovery that Neanderthals carried a variant, different from any known in modern humans, which is predicted to have been functional and cause pale skin [54]. Our results, we believe, strongly support the positive selection of *MC1R* variants, rather than simply relaxation of purifying selection in human groups that left high-UV zones. The positive correlation between strong lighter skin variants with latitude and decreasing UV exposure is likewise a strong indication of this. The lack of correlation between latitude/UV exposure between the ‘weaker’ variants (p.Val60Leu, p.Val92Met and p.Arg163Trp) may be due decreased selective pressure – most east Asian/east-central Asian populations live at a latitude comparable to southern Europe, so the selective pressure might be weaker and a smaller depigmentation effect might be sufficient. We also provide additional evidence for the convergent evolution of lighter skin within the Eurasian continent, as different variants with similar functions in a single, small gene, appear to have been selected in western and eastern populations.

**Fig 6.**
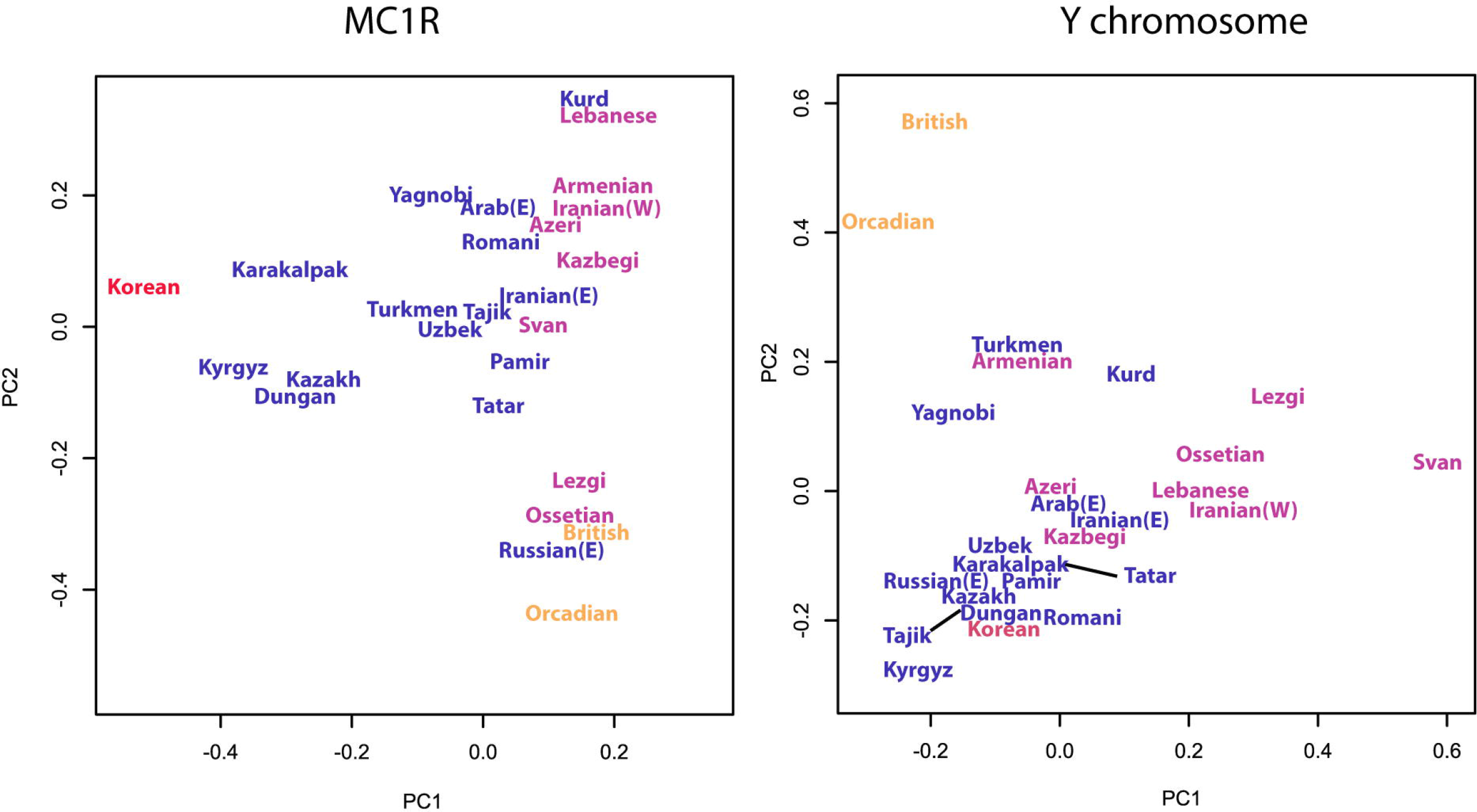
Principal components plots. Left: *MC1R* variants, where PC1 accounts for 91% of the variance, and PC2 4%. Right: Y chromosome data (from Wells et al. 2001 [3]), where PC1 accounts for 23% of the variance and PC2 19%. Populations are colour-coded; Europe - orange, West Asia - purple, Central Asia - blue, and East Asia - red.

## Conclusion

The high variability of mtDNA types, and congruent Y chromosome data, may be a source of human variation in both the east and the west of Eurasia. However, the role of subsequent migrations is difficult to evaluate. The lack of substantial correlation between language family groups and genetics beyond that determined by geography emphasizes the fact that genetic variation is usually stable compared to cultural variation, and that the association between the two is ephemeral. The *CCR5del32* distribution indicates its origin, but has no obvious association with any known epidemic, though selection with respect to viral pathogens remains a distinct possibility. The *MC1R* variant data provide another example of convergent selection for pale skin in the east and west. Large-scale genome-wide genetic data covering all of Eurasia, especially ancient DNA from a large span of dates, are needed to help clarify these issues and put our data into a wider context.

## Supporting Information

**S1 Fig. Sampling locations for individuals genotyped for the *CCR5* 32 bp deletion**. Locations are colored according to region, with Europe - orange, West Asia - purple, Central Asia - blue, South Asia - green, East Asia - red, and Southeast Asia - brown.

**S2 Fig. Language families in West and Central Asia – their geographic distributions and linguistic relationships.** a) The sampling locations of population samples speaking Iranian and Turkic languages, with each sample colour-coded according to language branch. b) Which languages correspond to the colour codes, and the linguistic relationship between the languages in question (upper panel for the Iranian language family, lower panel for the Turkic language family). c) PC plots of *MC1R* and Y chromosome with populations colour-coded according to language group rather than geographic location.

**S1 Appendix. Sample details and background.**

**S2 Appendix. Mitochondrial typing**

**S1 Table. Populations typed for *MC1R* variants.** ‘Population code’ is the unique identifier given to the sample set. Population codes with alphanumeric suffixes (e.g. Chinese4a & Chinese4b) are the same sampling, for which a different number of its constituent individuals has been typed for a given marker. ‘Geographic homeland’ is the area in which the ethnic/national group in question has its largest populations, ‘Sampling location’ is the exact place where the individual samples were collected, when known, ‘N’ is the number of individuals in the sample, and ‘Source’ details where the information was obtained.

**S2 Table. Populations typed for *CCR5del32* m.** ‘Population code’ is the unique identifier given to the sample set. Population codes with alphanumeric suffixes (e.g. Chinese4a & Chinese4b) are the same sampling, for which a different number of its constituent individuals has been typed for a given marker. ‘Geographic homeland’ is the area in which the ethnic/national group in question has its largest populations, ‘Sampling location’ is the exact place where the individual samples were collected, when known, ‘N’ is the number of individuals in the sample, and ‘Source’ details where the information was obtained.

**S3 Table. *MC1R* variant frequencies.** Count and frequency information for p.Val60Leu, p.Val92Met, p.Arg151Cys, p.Arg160Trp, p.Arg163Gln in populations from S1 Appendix.

**S4 Table. *CCR5del32* counts and frequencies.** Count and frequency information for *CCR5del32* in populations from S1 Appendix.

**S5 Table. Mitochondrial haplogroup frequencies.** The second column, ‘N’ gives the number of samples typed for each population group.

**S6-8 Table. F**_*ST*_ **values for all assessed variation.** S6 Table – Population pairwise F_*ST*_ values for the 32bp deletion in *CCR5*. The software F_*ST*_ estimation will occasionally output small negative values, these have been adjusted to 0. Higher F_*ST*_ values are highlighted; 0.05-0.1 in yellow, 0.1-0.15 in orange, and *>*0.15 in pink. Results significant at the 5% level or smaller are in bold; results significant at the 0.5% level or smaller are in larger font and bold.

S7 Table. Mitochondrial F_*ST*_ values. Population pairwise F_*ST*_ values for the mitochondrial HVRI sequences. The software F_*ST*_ estimation will occasionally output small negative values, these have been adjusted to 0. Higher F_*ST*_ values are highlighted; 0.02-0.04 in yellow, 0.04-0.06 in orange, and *>*0.06 in pink. Results significant at the 5% level or smaller are in bold; results significant at the 0.5% level or smaller are in larger font and bold.

ST8. MC1R F_*ST*_ values. This spreadsheet contains one sheet for each of the MC1R variants, each sheet giving the population pairwise F_*ST*_ value matrix. The software F_*ST*_ estimation will occasionally output small negative values, these have been adjusted to 0. High F_*ST*_ values are highlighted; 0.1-0.2 in yellow, 0.2-0.3 in orange, and *>*0.3 in pink. Results significant at the 5% level or smaller are in bold; results significant at the 0.5% level or smaller are in larger font and bold.

**S9 Table. F**_*ST*_ **values for combined *MC1R* variants.** Numbers in larger font and bold are considered significant. Values of 0.1-0.25 are shaded in green, 0.25-0.5 in yellow, and over 0.5 in orange.

## References

1. Krader L. Peoples of Central Asia. Bloomington: Indiana University Publications; 1966.

2. Bregel Y. An Historical Atlas of Central Asia. Leiden: Brill; 2003.

3. Wells RS, Yuldasheva N, Ruzibakiev R, Underhill PA, Evseeva I, Blue-Smith J, et al. The Eurasian heartland: a continental perspective on Y-chromosome diversity. Proc Natl Acad Sci U S A. 2001;98(18):10244–9.

4. Moore JP, Trkola A, Dragic T. Co-receptors for HIV-1 entry. Curr Opin Immunol. 1997;9(4):551–62.

5. Stephens JC, Reich DE, Goldstein DB, Shin HD, Smith MW, Carrington M, et al. Dating the origin of the CCR5-Delta32 AIDS-resistance allele by the coalescence of haplotypes. Am J Hum Genet. 1998;62(6):1507–15.

6. Agrawal L, Jin Q, Altenburg J, Meyer L, Tubiana R, Theodorou I, et al. CCR5Delta32 protein expression and stability are critical for resistance to human immunodeficiency virus type 1 in vivo. J Virol. 2007;81(15):8041–9.

7. Limborska SA, Balanovsky OP, Balanovskaya EV, Slominsky PA, Schadrina MI, Livshits LA, et al. Analysis of CCR5Delta32 geographic distribution and its correlation with some climatic and geographic factors. Hum Hered. 2002;53(1):49–54.

8. Valverde P, Healy E, Jackson I, Rees JL, Thody AJ. Variants of the melanocyte-stimulating hormone receptor gene are associated with red hair and fair skin in humans. Nat Genet. 1995;11(3):328–30.

9. Savage SA, Gerstenblith MR, Goldstein AM, Mirabello L, Fargnoli MC, Peris K, et al. Nucleotide diversity and population differentiation of the melanocortin 1 receptor gene, MC1R. BMC Genet. 2008;9:31.

10. Norton HL, Kittles RA, Parra E, McKeigue P, Mao X, Cheng K, et al. Genetic evidence for the convergent evolution of light skin in Europeans and East Asians. Mol Biol Evol. 2007;24(3):710–22.

11. Roberts DF, Kahlon DP. Environmental correlations of skin colour. Ann Hum Biol. 1976;3(1):11–22.

12. Jablonski NG, Chaplin G. The evolution of human skin coloration. J Hum Evol. 2000;39(1):57–106.

13. Winney B, Boumertit A, Day T, Davison D, Echeta C, Evseeva I, et al. People of the British Isles: preliminary analysis of genotypes and surnames in a UK control population. Eur J Hum Genet. 2012;20:203–210.

14. Comas D, Plaza S, Wells RS, Yuldaseva N, Lao O, Calafell F, et al. Admixture, migrations, and dispersals in Central Asia: evidence from maternal DNA lineages. Eur J Hum Genet. 2004;12(6):495–504.

15. Quintana-Murci L, Chaix R, Wells RS, Behar DM, Sayar H, Scozzari R, et al. Where west meets east: the complex mtDNA landscape of the southwest and Central Asian corridor. Am J Hum Genet. 2004;74(5):827–45.

16. Bartlett S, Straub J, Tonks S, Wells RS, Bodmer JG, Bodmer WF. Alkaline-mediated differential interaction (AMDI): a simple automatable single-nucleotide polymorphism assay. Proc Natl Acad Sci U S A. 2001;98(5):2694–7.

17. Excoffier L, Laval G, Schneider S. Arlequin ver 3.0: An integrated software package for population genetics data analysis. Evolutionary Bioniformatics Online. 2005;1:47–50.

18. Comas D, Calafell F, Mateu E, Perez-Lezaun A, Bosch E, Martinez-Arias R, et al. Trading genes along the silk road: mtDNA sequences and the origin of central Asian populations. Am J Hum Genet. 1998;63(6):1824–38.

19. Beaumont KA, Shekar SN, Newton RA, James MR, Stow JL, Duffy DL, et al. Receptor function, dominant negative activity and phenotype correlations for MC1R variant alleles. Hum Mol Genet. 2007;16(18):2249–60.

20. Frachetti MD. Migration Concepts in Central Eurasian Archaeology. Annu Rev Anthropol. 2011;40:191–212.

21. Zhong H, Shi H, Qi XB, Duan ZY, Tan PP, Jin L, et al. Extended Y-chromosome investigation suggests post-Glacial migrations of modern humans into East Asia via the northern route. Mol Biol Evol. 2010;.

22. Chaix R, Austerlitz F, Hegay T, Quintana-Murci L, Heyer E. Genetic traces of east-to-west human expansion waves in Eurasia. Am J Phys Anthropol. 2008;136(3):309–17.

23. Abdulla MA, Ahmed I, Assawamakin A, Bhak J, Brahmachari SK, Calacal GC, et al. Mapping human genetic diversity in Asia. Science. 2009;326(5959):1541–5.

24. Frenzel B, Pecsi M, Velichko AA. Atlas of Paleoclimates and Paleoenvironments of the Northern Hemisphere: Late Pleistocene-Holocene. Budapest: Hungarian Academy of Sciences; 1992.

25. Gonzalez-Ruiz M, Santos C, Jordana X, Simon M, Lalueza-Fox C, Gigli E, et al. Tracing the origin of the east-west population admixture in the Altai region (Central Asia). PLoS One. 2012;7(11):e48904.

26. Yunusbayev B, Metspalu M, Metspalu E, Valeev A, Litvinov S, Valiev R, et al. The Genetic Legacy of the Expansion of Turkic-Speaking Nomads across Eurasia. PLoS Genet. 2015;11(4):e1005068.

27. Hummel S, Schmidt D, Kremeyer B, Herrmann B, Oppermann M. Detection of the CCR5-Delta32 HIV resistance gene in Bronze Age skeletons. Genes Immun. 2005;6(4):371–4.

28. Sabeti PC, Walsh E, Schaffner SF, Varilly P, Fry B, Hutcheson HB, et al. The case for selection at CCR5-Delta32. PLoS Biol. 2005;3(11):e378.

29. Galvani AP, Novembre J. The evolutionary history of the CCR5-Delta32 HIV-resistance mutation. Microbes Infect. 2005;7(2):302–9.

30. Novembre J, Galvani AP, Slatkin M. The geographic spread of the CCR5 Delta32 HIV-resistance allele. PLoS Biol. 2005;3(11):e339.

31. Bollback JP, York TL, Nielsen R. Estimation of 2Nes from temporal allele frequency data. Genetics. 2008;179(1):497–502.

32. Novembre J, Han E. Human population structure and the adaptive response to pathogen-induced selection pressures. Philos Trans R Soc Lond B Biol Sci. 2011;367(1590):878–86.

33. Johansson A, Ingman M, Mack SJ, Erlich H, Gyllensten U. Genetic origin of the Swedish Sami inferred from HLA class I and class II allele frequencies. Eur J Hum Genet. 2008;16(11):1341–9.

34. Salmela E, Lappalainen T, Fransson I, Andersen PM, Dahlman-Wright K, Fiebig A, et al. Genome-wide analysis of single nucleotide polymorphisms uncovers population structure in Northern Europe. PLoS ONE. 2008;3(10):e3519.

35. Lappalainen T, Koivumaki S, Salmela E, Huoponen K, Sistonen P, Savontaus ML, et al. Regional differences among the Finns: a Y-chromosomal perspective. Gene. 2006;376(2):207–15.

36. Cavalli-Sforza LL, Menozzi P, Piazza A. Response. Science. 1993;261(5128):1508.

37. Huyghe JR, Fransen E, Hannula S, Van Laer L, Van Eyken E, Maki-Torkko E, et al. A genome-wide analysis of population structure in the Finnish Saami with implications for genetic association studies. Eur J Hum Genet. 2010;19(3):347–52.

38. Martinez-Cruz B, Vitalis R, Segurel L, Austerlitz F, Georges M, Thery S, et al. In the heartland of Eurasia: the multilocus genetic landscape of Central Asian populations. Eur J Hum Genet. 2011;19(2):216–23.

39. Flanagan N, Healy E, Ray A, Philips S, Todd C, Jackson IJ, et al. Pleiotropic effects of the melanocortin 1 receptor (MC1R) gene on human pigmentation. Hum Mol Genet. 2000;9(17):2531–7.

40. Bastiaens MT, ter Huurne JA, Kielich C, Gruis NA, Westendorp RG, Vermeer BJ, et al. Melanocortin-1 receptor gene variants determine the risk of nonmelanoma skin cancer independently of fair skin and red hair. Am J Hum Genet. 2001;68(4):884–94.

41. Sulem P, Gudbjartsson DF, Stacey SN, Helgason A, Rafnar T, Magnusson KP, et al. Genetic determinants of hair, eye and skin pigmentation in Europeans. Nat Genet. 2007;39(12):1443–52.

42. Jablonski NG. The Evolution of Human Skin and Skin Color. Annu Rev Anthropol. 2004;33:585–623.

43. John PR, Makova K, Li WH, Jenkins T, Ramsay M. DNA polymorphism and selection at the melanocortin-1 receptor gene in normally pigmented southern African individuals. Ann N Y Acad Sci. 2003;994:299–306.

44. Makova KD, Ramsay M, Jenkins T, Li WH. Human DNA sequence variation in a 6.6-kb region containing the melanocortin 1 receptor promoter. Genetics. 2001;158(3):1253–68.

45. Rogers A, Iltis D, Wooding S. Genetic Variation at the MC1R Locus and the Time since Loss of Human Body Hair. Curr Anthropol. 2004;45(1).

46. Harding RM, Healy E, Ray AJ, Ellis NS, Flanagan N, Todd C, et al. Evidence for variable selective pressures at MC1R. Am J Hum Genet. 2000;66(4):1351–61.

47. Rana BK, Hewett-Emmett D, Jin L, Chang BH, Sambuughin N, Lin M, et al. High polymorphism at the human melanocortin 1 receptor locus. Genetics. 1999;151(4):1547–57.

48. Lao O, de Gruijter JM, van Duijn K, Navarro A, Kayser M. Signatures of positive selection in genes associated with human skin pigmentation as revealed from analyses of single nucleotide polymorphisms. Ann Hum Genet. 2007;71(Pt 3):354–69.

49. Edwards M, Bigham A, Tan J, Li S, Gozdzik A, Ross K, et al. Association of the OCA2 polymorphism His615Arg with melanin content in east Asian populations: further evidence of convergent evolution of skin pigmentation. PLoS Genet. 2010;6(3):e1000867.

50. Cavalli-Sforza LL. Some old and new data on the genetics of human populations. Ala J Med Sci. 1966;3(4):376–81.

51. Mundy NI, Kelly J. Evolution of a pigmentation gene, the melanocortin-1 receptor, in primates. Am J Phys Anthropol. 2003;121(1):67–80.

52. Xue Y, Wang Q, Long Q, Ng BL, Swerdlow H, Burton J, et al. Human Y chromosome base-substitution mutation rate measured by direct sequencing in a deep-rooting pedigree. Curr Biol. 2009;19(17):1453–7.

53. Lewontin RC, Krakauer J. Distribution of gene frequency as a test of the theory of the selective neutrality of polymorphisms. Genetics. 1973;74(1):175–95.

54. Lalueza-Fox C, Rompler H, Caramelli D, Staubert C, Catalano G, Hughes D, et al. A melanocortin 1 receptor allele suggests varying pigmentation among Neanderthals. Science. 2007;318(5855):1453–5.

